# Joint tracking of active and passive motion in the mouse visual thalamus during natural behaviour

**DOI:** 10.1101/2025.11.03.686375

**Authors:** A.S. Ebrahimi, M.P. Hogan, F. Jin, F.P. Martial, Q. Huang, R. Joshi, K. Shirvanian, M. Burgess, K. Tang, Z. Montazeri, R.S. Petersen, R. Storchi

**Affiliations:** University of Manchester, School of Biological Sciences, Division of Neuroscience; University of Manchester, School of Engineering, Department of Computer Science

## Abstract

Most retinal motion is generated by the observer’s movements. These are often self-generated (active) but can also arise from external forces such as gravity (passive). Active movements modulate spontaneous and visually evoked activity throughout the visual system, providing neural circuits with information about behavioural state. By contrast, much less is known about how visual circuits represent passive movements. Critically, it remains unclear whether the retina-recipient thalamus tracks passive motion and, if so, whether this representation is separable from active motion.

To address these questions, we developed a paradigm that dissociates active and passive motion in freely moving mice and used it to examine their effects on neural activity. Mice actively explored an arena while intermittent floor tilts induced passive movement. By combining 3D reconstructions of animal pose and arena motion, we quantified active and passive linear and angular motion independently, even when they occurred simultaneously.

We found that neurons in the dorsal lateral geniculate nucleus (dLGN) and surrounding regions tracked combinations of active and passive motion signals both in the dark and during visual stimulation. Individual neurons differentially weighted active and passive motion signals. This diversity enabled active and passive motion signals to be linearly decoded and partially separated from population activity, while recurrent dynamics retained passive motion information over timescales of several seconds.

These findings identify a coding strategy through which early visual circuits integrate active and passive motion into an estimate of self-motion while preserving information about whether visual inputs arise from self-generated or externally imposed movements.

**Data availability:** Code, data and movies are available at https://doi.org/10.17605/OSF.IO/DRVFP

## Introduction

Although motion is mostly self-initiated (‘active’), animals also experience ‘passive’ motion imposed by features of the environment (e.g. slipping, falling, being pushed). Both active and passive motion alter the animal’s position and gaze direction and therefore shape the pattern of visual inputs reaching the retina. To correctly interpret this continuously changing input, visual circuits must track the animal’s motion state and integrate this information with ongoing visual processing.

It is now well established that motion state profoundly influences visual processing across species. Spontaneous activity is modulated by locomotion in Drosophila optic lobes [1], rodent visual thalamus and cortices [2-4], and peripheral visual field representations in primate V1 [5]. Motion state also strongly shapes visually evoked responses: swimming suppresses activity in the zebrafish optic tectum [6], locomotion exert well established effects on responses in rodent visual thalamus and cortex [3, 7-10], walking-cycle modulate responses in Drosophila visual circuits [11], and responses are suppressed before and during saccades, and subsequently enhanced, in the visual thalamus and cortex of cats and primates [12-14].

Thus, early visual circuits continuously track the animal’s motion state and integrate this information with incoming visual signals. Through this integration, motion-related signals can directly shape perception. For example, visual sensitivity in humans is modulated around the time of eye movements [15] and varies across the walking cycle during locomotion [16].

However, most previous studies have focused on active motion, which is generated by motor circuits and made available to other brain regions, including visual centres, through corollary discharges [17]. In contrast, passive motion is externally imposed, often unpredictable, and signalled primarily through vestibular and proprioceptive pathways [18, 19]. Only a few studies have examined its effects on early visual processing, primarily in visual cortex and under head-fixed conditions [20-22]. Consequently, it remains unknown whether subcortical regions such as the primary visual thalamus—the main gateway of visual information to cortex in mammals—also tracks passive motion during natural freely moving behaviour and, if so, whether active and passive motion are represented by separable neural signals or combined into a common estimate of motion state.

The most direct way to address these gaps consists in capturing the natural patterns of active and passive motion in freely moving animals since their complex interplay cannot be easily replicated in physically restrained (e.g. head-fixed) animals. To do so, we first developed a behavioural assay where freely animals freely explored a motorised arena which induced passive motion of the animal in the form of controlled tilts. By taking advantage of recent advances in tracking technology and 3D video reconstruction [2, 23, 24], we developed a method that dissected animal’s active and passive motion, even when these motion components occurred simultaneously.

We found that found that neurons in primary visual thalamus and neighbouring regions tracked both active and passive linear and angular head motion. Individual neurons differed in their sensitivity to such active and passive signals with most cells responding to multiple motion components but with different relative weightings. This diversity enabled both the tracking of overall self-motion and the partial separation of its active and passive origins from population activity through a simple linear decoder. Beyond such rapid responses, passive motion also engaged recurrent dynamics that generated temporally smooth representations of recent motion.

These results identify the primary visual thalamus as a tracker of motion state during natural behaviour, integrating active and passive motion while preserving information about their origin.

## Results

### Cells in visual thalamus jointly encode changes in light intensity and tilt events

To study how visually responsive cells respond to active and passive motion we first established a behavioural essay that enabled the animals to express their active motor repertoire while recreating conditions in which animals where passively moved. In this essay, naïve mice were placed in a small open field arena to record active exploration, and the arena was fitted with servo motors to produce floor tilts (20 x 20cm, **Figure 1A**). Arena floor and walls were transparent to allow visual stimulation from monitors placed around the arena but covered with a see-through metal grid that enabled animals to maintain a firm grip during floor tilts. To capture animal poses and arena position we employed 8 synchronised cameras (**Figure 1A**). Keypoints on both mouse and arena were tracked and 3D reconstructed by using custom developed software (**Figure 1B**; **Supplementary Movie 1**; see **Methods**). Left and right tilts were delivered from the same initial position in which the arena floor was parallel to the ground. After each of such perturbation the arena was maintained in tilted position for 1s, then tilted back to its initial position (**Figure 1C**). Tilts induced passive head/body rolls when the mouse was oriented parallel to the tilt axis, or head/body pitch, when oriented perpendicular to the tilt axis. Since animals thoroughly explored the arena in each experimental session, we were able to collect a large sample of roll and pitch events induced by the tilt (**Figure 1D**).

**Figure 1.**
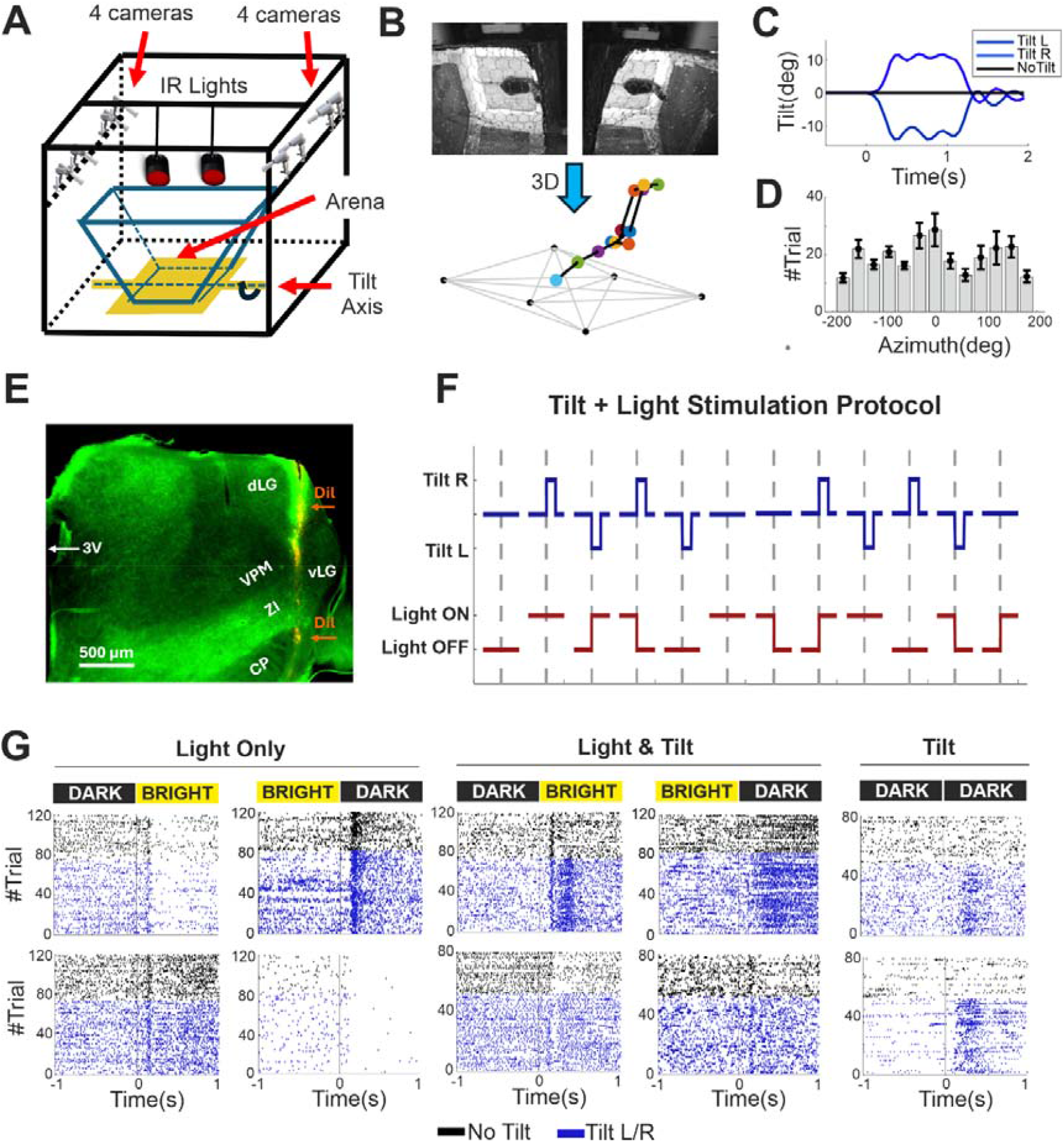
**A)** Schematic drawing of the behavioural recording set-up. **B)** Sample images from two cameras capturing the animal from different views, and the corresponding 3D reconstruction based on the full set of eight camera views. **C)** Rotation angles for left and right tilts and no tilt condition measured from 3D reconstructions of the behavioural videos. **D)** Histogram of the number of trials (Mean ± SEM; averaged across dataset of 9 animals) in which the animal head was oriented in specific directions (as shown on the x axis). **E)** Histological verification of the electrode placement. To visualise its position in the brain, the electrode was painted with fluorescent DIL marker before penetrating the brain. **F)** Schematic of the protocol for visual and tilt stimulation. Notice the 12 distinct trials encompassing all combinations of visual stimulation and tilts. **G)** Representative raster plots of individual units showing cells responding to light but not tilt events (Light Only), light and tilt events (Light & Tilt) and to tilt events in dark (Tilt). In a subset of trials (in which spikes are marked by black dots) the arena is maintained stationary. In the remaining trials (in which spikes are marked by blue dots) arena tilts are initiated at time 0. Arena illumination is provided by monitors mounted on the arena sides is turned from DARK to BRIGHT or BRIGHT to DARK at time 0.

To quantify neural responses, animals were implanted with Neuropixel 1.0 electrodes targeting primarily dLGN but also spanning more ventral thalamic and subthalamic nuclei (**Figure 1E**). To record responses to visual stimulation, three types of visual stimuli were delivered from monitors mounted around the arena: a step from dark to bright light (DARK-BRIGHT), a step from bright light to dark (BRIGHT-DARK) and constant bright light (BRIGHT-BRIGHT). To isolate responses to tilt events from responses to visual stimulation, additional experimental trials were recorded under constant darkness (DARK-DARK). The visual stimuli were combined with two tilt stimuli comprising a left and a right left tilt. To isolate responses to visual stimuli from tilt events we recorded additional experimental trials without a tilt. Overall, by combining visual and tilt stimulation we generated 12 distinct types of experimental trials (4 visual x 3 tilt, see protocol schematic in **Figure 1F**). The different visual stimuli were interleaved while the occurrence and direction of tilts followed a pseudorandom sequence to reduce the animal’s ability to predict them. For each animal, we recorded 400 trials (∼33 per condition). Statistical tests (see **Methods**) were used to identify cells that were only responsive to visual stimulation, cells whose visual responses were affected by tilts and cells that responded to tilts in the dark (**Figure 1G**).

We first analysed neuronal responses to light stimulation without arena tilts (n=498). A summary of light responsive units across animals and brain regions in our dataset is provided in **Supplementary Table 1**. Within this dataset, visual stimulation evoked a variety of response types and a clustering algorithm, spanning a wide range of clustering solutions (**Methods**), identified 5 main classes (**Figure 2A, C, left panels; Figure 2B, D, orange lines**). We then measured visual responses from the same cells but during tilt events. We found that these events substantially affected firing rates (**Figure 2A, C, mid panels; Figure 2B, D, blue lines**). At single cell level, this effect was significant in a sizable fraction of cells, both in dLGN (41%, 119/290) and neighbouring regions (VPM: 30%, 20/67; ZI: 53%, 44/82; CP: 56%, 33/59). Moreover, the tilts affected all classes of light responses (p < 0.00001 for DARK-BRIGHT and p < 0.01 for BRIGHT-DARK across all 5 classes; sign-test, n = 150, 150, 87, 37, 74) and the net effect was a larger increase in firing rates compared with trials in which light stimulation was not paired with a tilt (**Figure 2E, F**, **G, H**). Individual classes differed in the amount to which firing rate was affected for responses to BRIGHT-DARK stimulation (p = 0.002, χ^2^ = 16.92, n = 498) but not for responses to DARK-BRIGHT stimulation (p = 0.12, χ^2^ = 7.33, n = 498). These results indicate that tilt events affect visual responses cells in visual thalamus and neighbouring regions, these effects are widely presents across diverse functional cell types while their size significantly differs across types.

**Figure 2.**
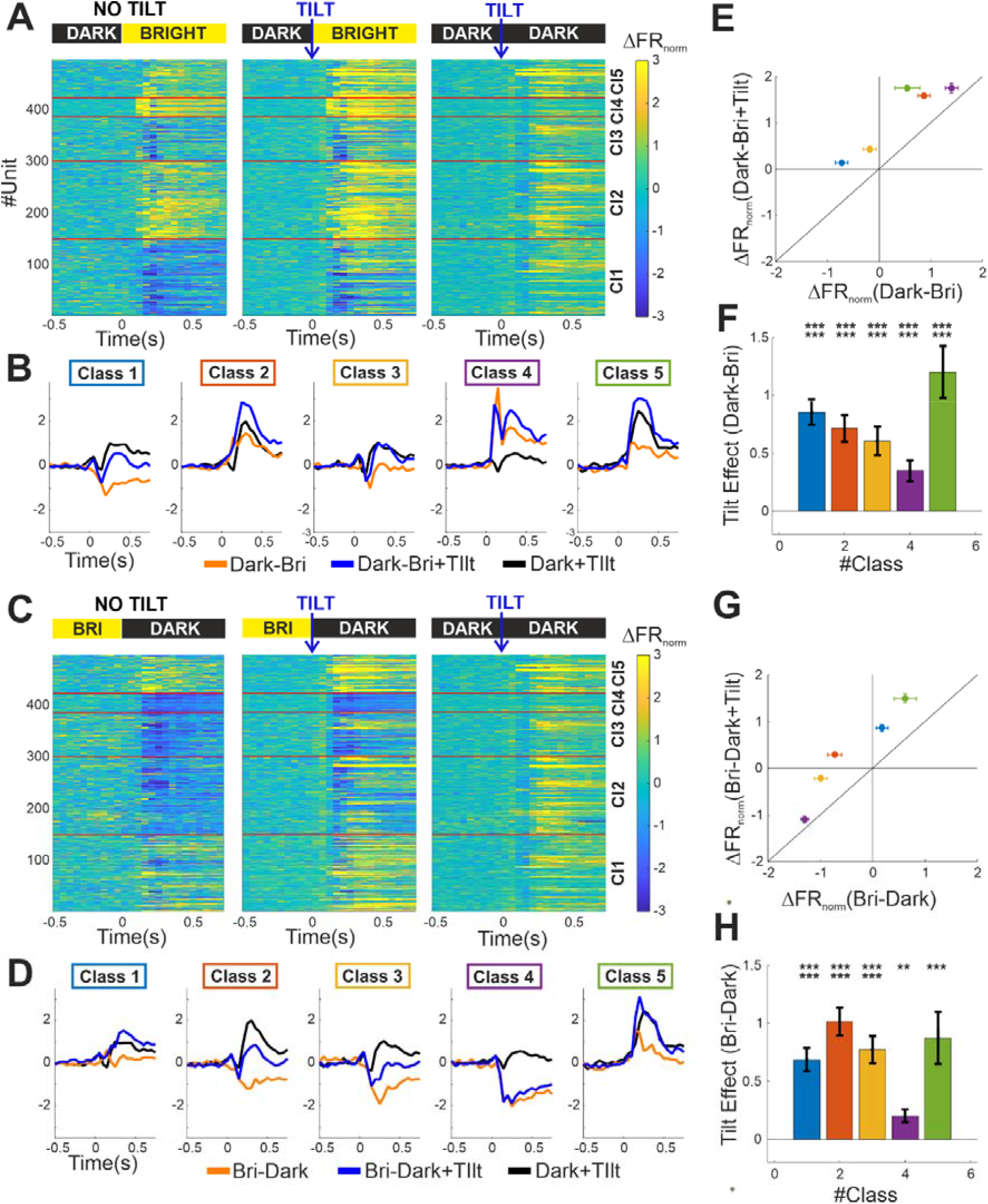
**A)** The left panel shows normalised PSTHs of single unit visual responses (DARK-BRIGHT) when the platform was kept stationary. Units are ordered along the y-axis according to a clustering analysis (see **Methods**). The transition between clusters is shown by red lines. The middle panel shows the same units during the same light stimulation but recorded in trials in which arena tilts were initiated at time 0. The right panel shows the same units recorded in the dark during arena tilts. **B)** Average PSTH responses across the five classes of visual responses determined by our cluster analyses. Averages are separately calculated during DARK-BRIGHT visual stimulation when the platform was kept stationary (orange), during the same visual stimulation when the platform was tilted (blue) and during tilt stimulation in the dark (black). **C)** Same as panel A but here we recorded BRIGHT-DARK visual stimulation. The right panel is replicated from panel A for clarity of visualisation. **D)** Same as panel B but here we recorded BRIGHT-DARK visual stimulation. Black lines are replicated from panel A. **E)** Population responses (Mean ± SEM) to visual DARK-BRIGHT visual stimulation when the platform was kept stationary (x-axis) vs responses to the same visual stimulation during tilt events (y-axis). Cell populations are divided according to the class of visual response (same colour convention used in panel **B. F)** Changes in visual responses, denoted as Tilt Effect (Mean ± SEM), obtained by subtracting visual responses during tilt stimulation from visual responses in absence of tilt. **G)** Same as panel E but here we recorded BRIGHT-DARK responses. **H)** Same as panel **F** but here we recorded BRIGHT-DARK responses.

If the tilt effect was visually driven (e.g. by fast movements of the visual scene projected on the animal’s retina) we would expect it to disappear in the dark. To test for this possibility, we delivered an additional pseudorandomised sequence of tilts (n=100 trials, DARK-DARK condition) after turning off all monitors and all the other light sources in the room. We found that a comparable fraction of units (dLGN: 34%, 99/290; VPM: 30%, 20/67; ZI: 45%, 37/82; CP: 63%, 37/59) responded to tilts in the dark (**Figure 2A, C, right panels; Figure 2C, D, black lines**). Moreover, a sizable fraction of units whose light responses were modified by the tilt also responded to tilt in the dark (dLGN: 19%, 55/290; VPM: 13%, 9/67; ZI: 33%, 27/82; CP: 49%, 29/59). These results indicate that the effect of tilt events in visual thalamus and neighbouring regions is largely nonvisual in nature.

### Responses to tilt events are robust to changes in baseline arousal levels in awake animals

In principle, neuronal responses to tilt could be fully explained by a change from a low to a high arousal state following tilt onset. To test whether tilt-evoked responses are robust across different baseline levels of arousal, we experimentally elevated arousal prior to tilt onset. To do this, we modified our original experimental protocol by delivering a loud sound (1046.502 Hz at 102 dB; duration = 0.5s) that reliably cued the occurrence of a tilt 1.5s later. No sound was delivered in trials in which the platform was kept stationary. To further increase animal’s arousal state, the platform was also vibrated throughout the 2s epoch preceding the tilt (see protocol schematic in **Figure 3A**).

**Figure 3.**
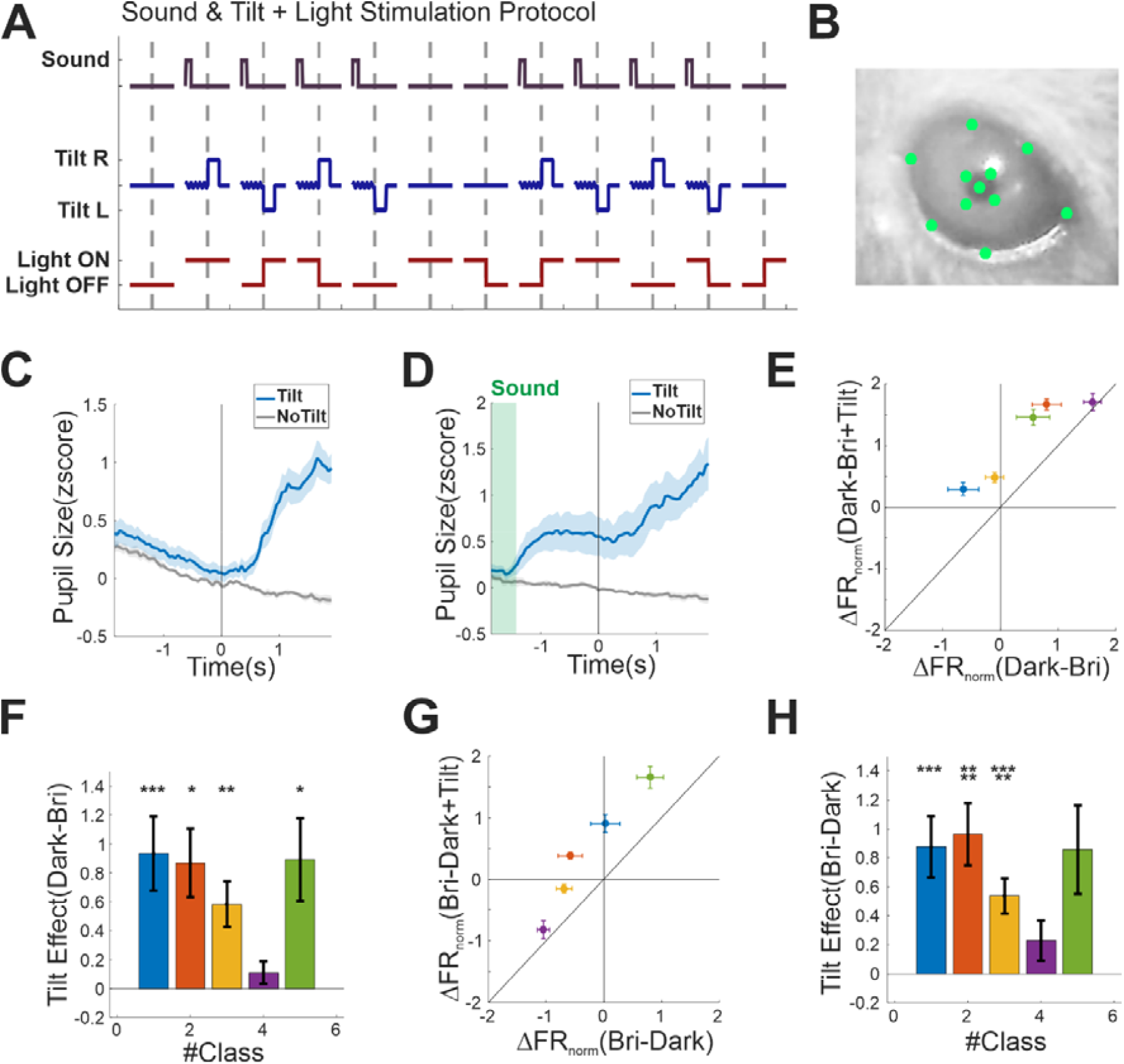
**A)** Schematic of the modified protocol. Here a sound precedes the onset of the tilt and in these trials the platform is vibrated before the tilt onset. In trials in which the platform is kept stationary no sound or vibrations are delivered. **B)** Example image of the mouse eye obtained during our experiments. Labelling of the eye landmarks used to track pupil size and stabilise the image are shown in green. **C)** Normalised pupil size during trials in which the platform was kept stationary (grey) or tilted (blue) for the original protocols described in Figure 1F. **D)** Normalised pupil size measures for the modified protocol shown in panel **A. E-H)** Same as **Figure 2E, F, G**, H but here we report data collected during the modified protocol in panel **A**.

We compared the effects of the two protocols on arousal state by running separate eye tracking experiments (n=4 animal) in which we measured changes in pupil diameter before and after the tilt events by keeping arena illumination constant to discount light-driven effects (**Supplementary Movie 2, Figure 3B**). In the original experiment robust changes in pupil diameter occurred only after the tilt onset (**Figure 3C**). Instead, in the modified protocol, changes in pupil diameter triggered by sound and vibrations started ∼1.5s before the tilt (**Figure 3D**). Nevertheless, pupil diameter further increased after the tilt onset, indicating that tilt-evoked pupil responses persist despite elevated pre-tilt arousal levels.

We then recorded animals neuronal activity under this modified protocol. We found that tilt events affected visual responses to an extent comparable with the original experiment and across most classes of light responsive cells (**Figure 3E-H**).

These results indicate that tilt-evoked responses are preserved across elevated arousal states and cannot be explained solely by a transition from a low to a high arousal state. However, tilt responses were strongly diminished in deeply anaesthetised animals (1.5mg/kg urethane inducing a surgical plane of anaesthesia, see **Supplementary Results**), indicating that a baseline awake brain state is required for their full expression.

### Three-dimensional reconstruction enables to dissect active and passive motion signals

We next developed a method to separate active and passive motion signals (see **Methods** for details) during tilt events by focussing on animal’s linear and angular head movements.

Firstly, we applied 3D keypoints reconstruction to capture head position and orientation across consecutive frames (denoted *X*(*i* − 1 ) and *X*(*i*) in **Figure 4A, OBSERVED MOTION**) in respect to the centre of the arena at rest (denoted *X*_0_). To discriminate active and passive motion signals we decomposed these initial tracking data along two distinct reference frames (see **Methods** for a detailed derivation). The first reference frame, defined as platform (**Figure 4A, REFERENCE FRAMES**) was positioned in the arena’s centre as before but locked to platform motion so that head’s position and orientation on the arena were invariant to arena’s tilt. The second reference frame, defined as fixed (**Figure 4A, REFERENCE FRAMES**) mapped the transformations between centre at rest and centre during tilt events. In this way animal’s position (denoted *X*_*a*_ could be expressed in respect to the platform’s frame, by computing rotation and translation (denoted *R*_*a*_ and *T*_*a*_)from platform’s centre, and in respect to the fixed frame (denoted *X*), by computing the transformation between frames (denoted *R*_*p*_ and *T*_*p*_)

**Figure 4.**
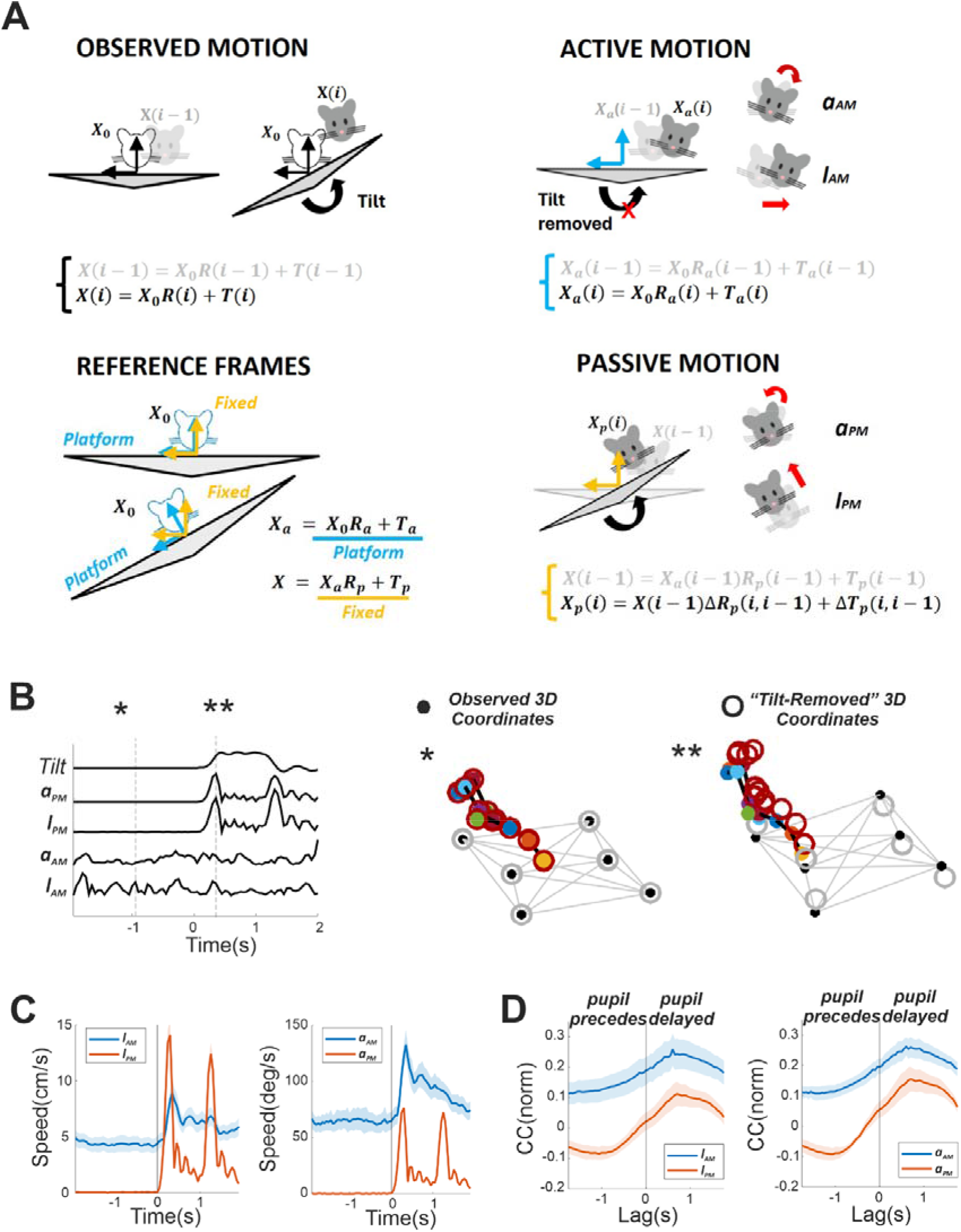
**A)** Schematic illustrating how active and passive motion signals are separated from 3D tracking data. Animal head positions in consecutive images (coloured heads) are initially estimated according to their rotation and translation in respect to a fixed reference (denoted by the empty head; OBSERVED MOTION). The initial tracking data are decomposed into rotations and translations in respect to a continuously updated coordinate system aligned with the platform (denoted as **Platform**) and rotations and translations between such reference and a fixed reference (denoted as Fixed, **REFERENCE FRAMES**). Active motion is measured by correcting head positions by removing the transformation between reference frames (**Tilt Removed**). The overall motion between consecutive images is then separated into linear and angular motion (ACTIVE MOTION). Passive motion is measured by “freezing” the animal pose on the platform (denoted by *X*(*i* - 1)) and propagating its position to the following sample according to movement of the platform (denoted *X*_*p*_(*i*), **PASSIVE MOTION**). **B)** An example trial showing normalised readouts of tilt angle, angular passive head speed (**a**_**PM**_), linear passive head speed (**l**_**PM**_), angular active head speed (**a**_**AM**_) and linear active head speed (**l**_**AM**_). On the right, we show two mouse poses recorded at two different time points (indicated by asterisks) within the trial. The 3D coordinates reconstructed from the video are indicated by coloured dots (Tracked 3D Coordinates) while coordinates used for measuring active motion signals, in which reference transformations were removed (“Tilt Removed”), are indicated by empty circles. **C)** Linear and angular head speed for active (blue error bars) and passive (red error bars) motion. **D)** Normalised cross correlation between active and passive motion signals (respectively blue and red error bars) and timeseries of pupil size.

To estimate active motion, we first removed the component of pose change *R*_*p*_ and *T*_*p*_ that was induced by platform tilt by expressing the animal’s pose in the platform’s frame. Active linear and angular motion (respectively *l*_*AM*_ and *a*_*AM*_ ) were then derived from the rotation and translation between consecutive poses (denoted as *X*_*a*_(*i*) and *X*_*a*_(*i* − 1 ) by calculating the magnitude of angular and linear displacements (**Figure 4A, ACTIVE MOTION**).

Passive motion was estimated by propagating the animal’s pose from time point *i*−*1* using the measured platform rotation Δ*R*_*p*_(*i, i* − 1 ) and translation Δ*T*_*p*_(*i, i* − 1 ) between consecutive time points, i.e. *X*_*p*_(*i*) = *X*(*i* − 1 ) Δ*R*_*p*_(*i, i* − 1 ) + Δ*T*_*p*_(*i, i* − 1 ) (**Figure 4A, PASSIVE MOTION**). The magnitude of passive angular and linear displacements (respectively *l*_*PM*_ and *l*_*PM*_ ) were then derived from such rotation and translation.

An illustration of the derived motion signals (*l*_*AM*_, *a*_*AM*_,*l*_*PM*_, *a*_*PM*_), together with a comparison between poses measured in a fixed external and a platform aligned reference frame is provided in **Figure 4B** and **Supplementary Movie 3**. Across our dataset we found that active motion signals l_AM_ and a_AM_ peaked soon after the onset of the tilt, remained elevated while the arena was in tilted position, then quickly decreased after the arena tilted back to its initial position (**Figure 4C, blue**). Passive head motion signals measuring *l*_*PM*_ and *a*_*PM*_ more closely followed tilt dynamics showing two distinct peaks, which corresponded to the tilt away from, and back to, the initial arena position (**Figure 4C, red**). A cross-correlation analysis revealed that both active and passive motion signals had distinct temporal dynamics compared with changes in pupil size, with both signals substantially anticipating pupil responses (**Figure 4D**).

### Cells in visual thalamus track heterogeneous combinations of active and passive motion signals

We next set-out to determine the relative extent to which active light responsive units in dLGN and surrounding regions track active and passive motion during tilt events. To address this question, we estimated predictive models of single unit firing activity based on motion signals and used these models to determine which motion signals were tracked by individual cells.

A key challenge in this analysis is that different motion signals are correlated. For example, as shown in **Figure 4C**, passive motion caused by tilts is often accompanied by increased active motion. This could lead to incorrect estimates of the dimensionality of the model (i.e. how many motion signals a cell responds to) and to the incorrect identification of which variables a neuron is responding to.

To address this problem, we first decorrelated components of active and passive motion signals by using two well established methods: Independent Component Analysis (ICA) and Principal Component Analysis (**Figure 5A**). We applied separately these methods to motion signals pooled across all animals implanted with brain electrodes and used for this study (∼5 hours recordings across 9 animals). To also capture temporal correlations, each original signal (*l*_*PM*_, *a*_*PM*_, *l*_*AM*_, *a*_*AM*_) was stacked across two consecutive time points (each of 50ms duration). Overall, the first six principal components (PCs) explained 97.42% of the variance in the data. An ICA model with the same dimensionality achieved a comparable reconstruction accuracy, accounting for 97.41% of the total variance. The PCs captured mixed combinations of active and passive movements with the 1^st^ PC computing a temporally weighted average of these signals – i.e. self-motion – while the 2^nd^ PC computing their difference (**Figure 5A**, *W*_*PC*_). The independent components (ICs) returned a sparser representation with the first two ICs computing temporally weighted average of active and passive motion signals respectively (**Figure 5A, *W***_***IC***_). Later PCs and ICs computed similar features encompassing the difference between linear and angular speeds (3^rd^ and 4^th^ components) and the accelerations in active and passive motion (respectively 5^th^ and 6^th^ components).

**Figure 5.**
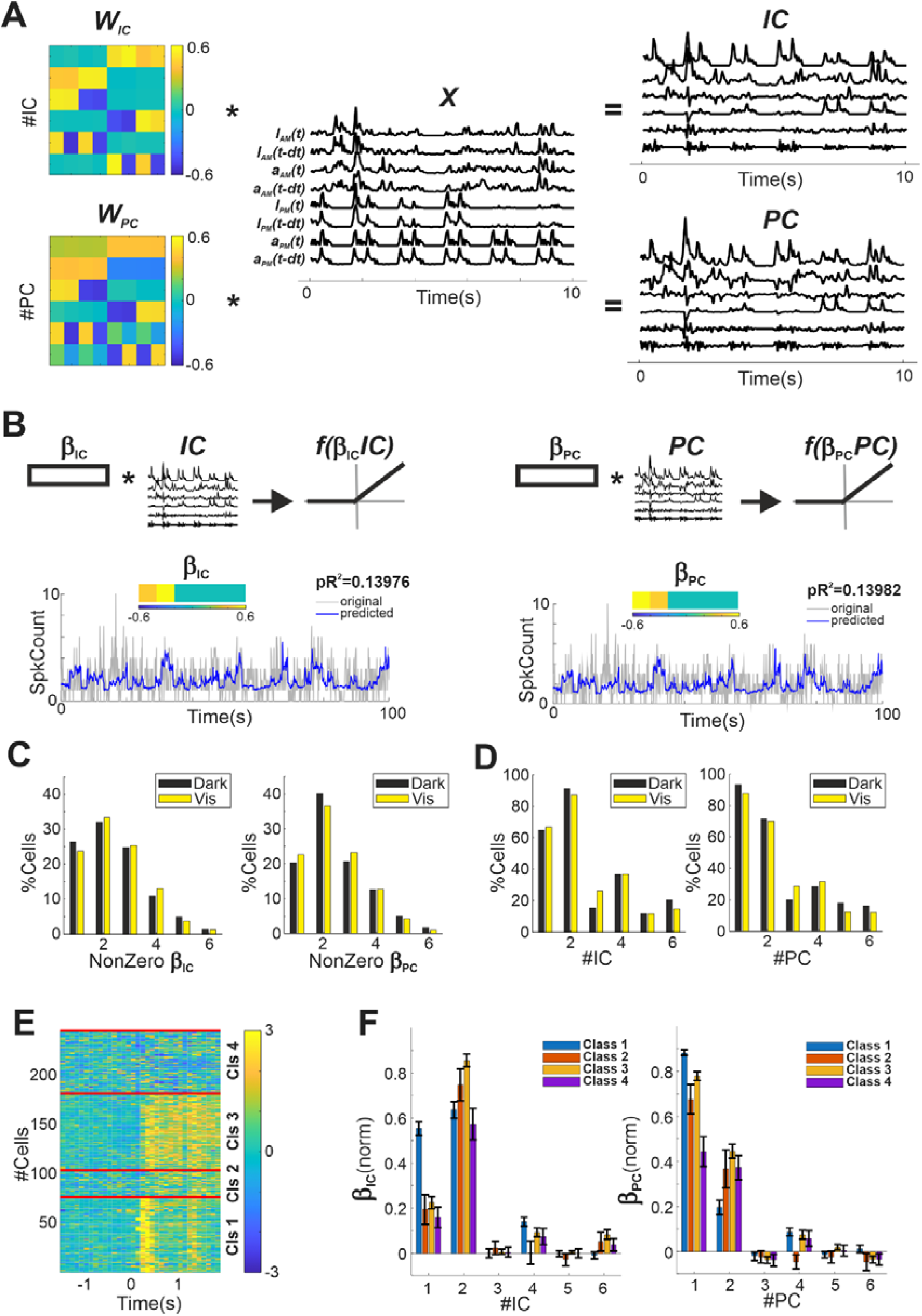
**A)** Independent and Principal Component Analyses of motion signals. The original signals (denoted X) are multiplied by the unmixing matrix for ICA (denoted *W*_*JC*_ ) and the loading matrix for PCA (denoted *W*_*PC*_) to obtain *IC* and *PC* matrices. These maximise statistical independence (*IC*) and enforce orthogonality (*PC*) among covariate signals making neural encoding models more easily interpretable. **B)** Schematic of encoding models for a representative single cell based on *IC* and *PC* signals. Cross-validated prediction accuracy as pseudo-R^2^ is reported next to panel legends. **C)** Percentage of cells, among those encoding motion signals based on Lasso variable selection (see Methods), encoding one or more covariate (denoted as NonZero *β*_*IC*_ and NonZero *β*_*PC*_). **D)** Percentage of cell encoding each of the six *IC* and *PC* covariates. **E)** A clustering analysis identifies four distinct classes of neurons responding to motion signals. **F)** Average weighting of different motion signals differ across the distinct classes shown in panel **E** (Mean ± SEM; coefficients scaled to have a unitary norm).

We employed the decorrelated motion signals ( ***IC*** and ***PC*** in **Figure 5A**) to fit Poisson Generalised Linear Models (pGLMs) of single cells activity (**Figure 5B**). The coefficients of pGLMs (***βIC*** and ***β***_***PC***_ in **Figure 5B**) were then used to determine which motion signals individual cells responded to. To improve pGLMs predictive ability, these models were augmented with additional covariates measuring animal orientation within the arena, visual stimuli (when present) and motion-independent fluctuations in population activity (see **Methods** for details). Since individual cells might not respond to any motion signal or respond only to a one or few of them, we first used lasso regularisation to ensure signals not useful for predictions were weighted at zero (see **Methods**). We then used the remaining nonzero signals to fit a non-penalised regression (see example regression in **Figure 5B**).

We found that both in data recorded in darkness and during visual stimulation a sizable fraction of cells encoded at least one decorrelated motion signal by using both ICs and PCs (DARK: 50%, 247/498; VIS: 67%, 333/498). Prediction performances in darkness and under light stimulation (DARK IC: 0.09 ± 0.080 SD, n = 247 cells; VIS IC: 0.09 ± 0.079 SD, n = 333 cells) were not significantly different between IC and PC decorrelated signals (DARK: p = 0.074; VIS: p = 0.330; t-tests). Among these cells, most responded to multidimensional combinations of two or more than motion components (**Figure 5C**; DARK: 74% for ICs and 80% PCs; VIS: 76% for ICs and 77% for PCs). The first two motion components for both ICs and PCs were tracked by the majority of these cells while the remaining ones only by smaller fractions (**Figure 5D**).

Moreover, individual cells tracked diverse combinations of motion signals. A clustering analysis, applied to neurons recorded in darkness to isolate the effect of motion, showed four different classes of cells (**Figure 5E**). These classes assigned different weights to individual motion signals (**Figure 5F**). For example, **Class 1** assigned similar weights to the first two ICs while **Class 3** primarily encoded the 2^nd^ IC (**Figure 5F**).

Together, these results indicate that motion is encoded in a distributed manner across the visual thalamus and surrounding regions, with individual neurons tracking different combinations of active and passive motion signals.

### Active and passive motion components are linearly decodable and partially separable from population activity

The distributed encoding of active and passive motion at the single-cell level in dLGN and surrounding regions raises the question of how these signals can be readout and to what extent they can be separated at the population level. Linear decoding provides a biologically plausible mechanism for such readout, as neurons can combine synaptic inputs through weighted summation. Thus, we asked whether a linear decoder could recover and separate active and passive motion components from population activity (**Figure 6A**). For this analysis, we selected recordings in which we simultaneously acquired at least 20 cells that responded to both visual stimulation and movements (n = 5 animals).

**Figure 6.**
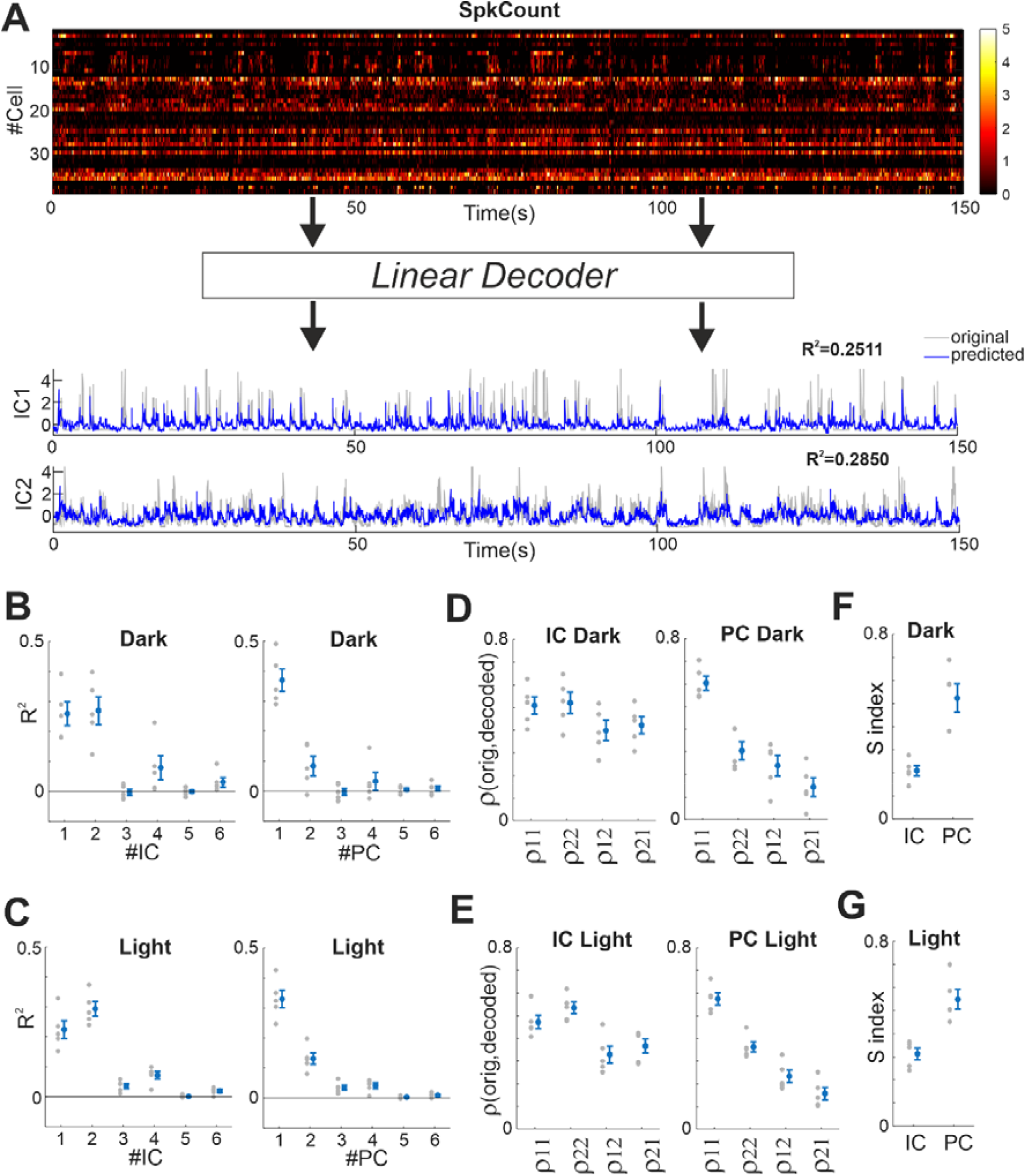
**A)** Representative population recording in dark during spontaneous exploration and tilt events (top panel, **SpkCount**). Motion signals (IC1 and IC2) are predicted with a linear decoder trained with ridge regression (bottom panel). **B)** Prediction accuracy (as **R**^**2**^) for IC and PC signals in the dark. The two first signals (IC1,IC2; PC1, PC2) are the ones predicted with highest accuracy across animals (n=5). **C)** Same as panel B but here animals are subjected to visual stimulation. **D**,**E)** Pearson’s correlation between the first two original and predicted motion signals for *IC* and *PC* in the dark. Nonzero correlation between an original signal and a different predicted signal (*ρ*_12_, *ρ*_21_) indicate some level of crosstalk in dark **(D)** and under light stimulation **(E). F**,**G)** Different motion signals are partially separable, as indicated by positive S index, both in dark (F) and under light stimulation **(G)**.

In the dark, the first two ICs and PCs could be decoded significantly above chance (**Figure 6B**, p=0.0034, 0.0050 for 1^st^ and 2^nd^ ICs; p=0.0010, 0.0217 for 1^st^ and 2^nd^ PCs; t-test on n=5 animals). Similar results were obtained under light stimulation (**Figure 6C**, p=0.0019, 0.0053 for 1^st^ and 2^nd^ ICs; p=0.0005, 0.0017 for 1^st^ and 2^nd^ PCs; t-tests on n=5 animals).

We then examined the extent to which the decoded signals were separable, focussing on the first two ICs and PCs. If two motion components are separable, a decoded component should correlate more strongly with its matching component (e.g. decoded IC1 versus original IC1) than with the non-matching component (e.g. decoded IC1 versus original IC2). This was the case for both PCs and ICs in the dark (**Figure 6D**, p<0.0005 for ICs; p<0.005 for PCs; t-tests on n=5 animals) and during light stimulation (**Figure 6E**, p<0.0005 for ICs; p<0.0001 for PCs; t-tests on n=5 animals). However, crosstalk between components was present in both conditions since both unmatched components also exhibited positive correlations (**Figure 6D**, p<0.001 & p<0.05 for ICs & PCs; **Figure 6E**, p<0.001 & p<0.005 for ICs & PCs). To quantify separability, we defined a separations index

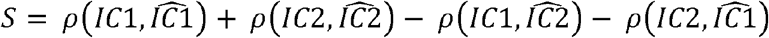

where positive values indicate stronger correlations between matching than non-matching components. We found *S* to be positive across ICs and PCs both in the dark ( **Figure 6F**, p=0.0006 for ICs, p=0.0010 for PCs, t-tests on n=5 animals) and under light stimulation (**Figure 6G**, p=0.0003 for ICs, p=0.0020 for PCs, t-tests on n=5 animals).

Together, these results indicate that a simple linear readout can be used decode active and passive motion signals from population activity in visual thalamus and the decoded signals are partially separable irrespectively of the lighting environment.

### Passive motion signals engage recurrent dynamics that maintain temporally extended representations

While our results demonstrate that dLGN and surrounding regions track passive motion through rapid responses, passive motion signals could in principle also engage longer-lasting recurrent dynamics. If present, such dynamics would allow neural circuits to retain information about recent motion history and maintain temporally smooth representations of self-motion.

Thus, we sought to dissociate the direct effects of passive motion signals, sampled in 50 ms time bins, from delayed responses mediated by recurrent dynamics. To achieve this, we applied the recently developed Input Preferential Subspace Identification (IPSID) framework [25] (see **Methods** for details). IPSID estimates a latent dynamical system containing dimensions that jointly drive neural activity and behaviour, alongside dimensions specific to either domain. Within this framework, passive motion signals were modelled as external inputs, allowing us to quantify both their direct impact on neural responses and their indirect influence through recurrent latent dynamics.

Simulations of IPSID models fitted to our datasets showed that the IPSID model captured average behavioural fluctuations measured by active motion signals (*l*_AM_ : ρ = 0.4641 ± 0.0611 SD; *a*_AM_ : ρ = 0.4464 ± 0.0811 SD; n=5 animals) and fluctuations in neural activity (ρ = 0.5611 ± 0.0734; **Figure 7A**) in darkness. Similar performance was observed under light stimulation (*l*_*AM*_ : : ρ = 0. 4520 ± 0. 0547 SD; *a*_*AM*_ ρ = 0. 4753 ± 0. 0797 SD; neural activity: ρ = 0. 4409 ± 0. 1335 SD; n=5 animals; **Figure 7E**). Having established that the model captured the measured behavioural and neuronal activity, we selectively ablated the direct and recurrent pathways to quantify their respective contributions to neural responses. We found that both components contributed to neural activity. In the dark, recurrent dynamics consistently accounted for a larger fraction of the explained variance across all animals (p = 0.0277, t-test on n = 5 animals; **Figure 7B, C**). During light stimulation, where recurrent dynamics also included those associated with visual stimuli, the variance explained by direct and recurrent dynamics was comparable (p = 0.3957, t-test on n = 5 animals; **Figure 7F, G**).

**Figure 7.**
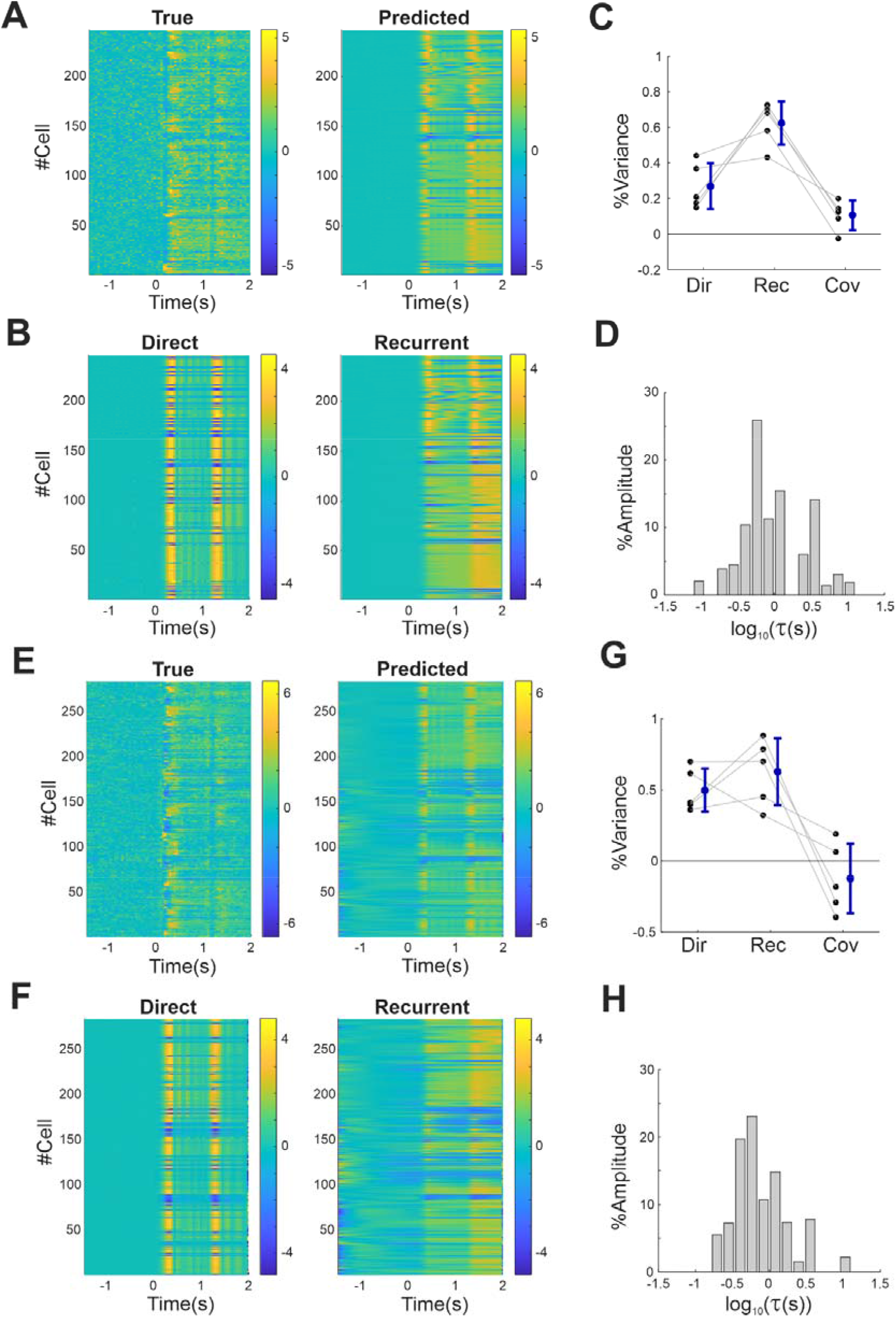
**A)** Z-scored PSTHs from neurons recorded in the dark (**True**) and predictions based on the cross-validated IPSID model (**Predicted**). **B)** Separation of neural activity driven directly by passive motion inputs (**Direct**) and indirectly through the recurrent latent dynamics engaged by those inputs (**Recurrent**). **C)** Variance decomposition of neural activity attributable to direct passive motion inputs (**Dir**) and recurrent dynamics (**Rec**). The remaining variance is explained by the covariance between these components (**Cov**). **D)** Distribution of neural loadings across latent dynamical timescales. Bars show the percentage amplitude of eigenvector projections onto neural activity as a function of the characteristic time constant of each eigenmode (x-axis, log scale, see **Methods** for details). **E-G)** Same as panels (**A-D**) but here neurons were recorded during visual stimulation.

We next estimated the timescales over which recurrent dynamics influence neural activity (see **Methods**). To do so, we determined the eigenvectors of the latent dynamical system and projected them onto the recorded neural population using the observation matrix estimated by IPSID. Since each eigenvector is associated with a characteristic time constant, this analysis allowed us to determine how strongly neurons represented latent dynamics operating on different timescales. At the population level, neural activity reflected a broad range of temporal dynamics, spanning hundreds of milliseconds to several seconds both in the dark (**Figure 7D**) and under light stimulation (**Figure 7H**).

Together, these results indicate that early visual circuits, beyond tracking rapid changes in passive motion signals, can also maintain temporally extended representations of such signals.

## Discussion

During natural behaviour, the patterns of light reaching the eye are shaped both by movements of the observer and by changes in the external world. To interpret this continuously changing input, visual circuits must determine whether changes in visual input arises from self-motion or external events and, when it arises from self-motion, whether that motion was self-generated (active movements) or externally imposed (passive movements). Here we found that during freely moving exploration dLGN and neighbouring regions track both active and passive motion signals. We further show that responses to passive motion signals are partially separable from those associated with active motion and are propagated in time through recurrent dynamics.

The thalamocortical pathway is considered the main route for visual perception. Along this pathway, motion processing has been primarily studied in primary visual cortex and at later cortical stages. Thus, responses to active eye, head and body movements, and to local optic flow have been well described in primary visual cortex (mouse [8, 10, 26, 27]; primates [14, 28, 29]) while higher cortical areas have been shown to respond to global optic flow (primates [30, 31]; mice [32, 33]). These visual regions are also known to integrate visual inputs with vestibular and motor signals [20-22, 34, 35], which could contribute to distinguishing self-motion from externally driven motion.

However, there is substantial evidence across mammalian species that signals related to the observer’s own movements are already present at subcortical stages and at the level of the dorsal lateral geniculate nucleus (dLGN). Both in dark and under light stimulation, neuronal activity around saccadic eye movements is suppressed prior to and enhanced following the saccade in this region [12, 14]. Locomotion and up/down head pitch have also been shown to modulate spontaneous and visually evoked activity in rodents dLGN [2, 3, 7].

The effects of passive motion on dLGN are less understood. Nevertheless, several lines of evidence suggest that dLGN could track passive motion. A study in pharmacologically immobilised cats found a large proportion of dLGN neurons responding to onset and offset of horizontal head rotations [36]. Studies in anaesthetised mice and cats have also shown that electrical and optogenetic stimulation of vestibular regions affect spontaneous and visual evoked activity in visual thalamus [37-39].

In spite of this body of work, a demonstration that dLGN can effectively track passive motion during natural behaviour, and how these signals were integrated with internal dynamics, was lacking. It was also unknown whether passive motion signals, if present, were integrated with or represented separately from active motion signals.

Our study addresses these gaps, substantially extending our understanding of how early visual circuits process self-motion during natural behaviours. By developing a new freely moving behavioural assay and by building on recent progress in 3D tracking techniques [2, 23, 24] we were able to separate linear and angular active and passive motion under natural, unrestrained conditions. Our results indicate that neurons in dLGN and neighbouring regions encode a heterogeneous representation of active and passive head motion. Through this diversity, passive motion signals can be integrated with active motion signals to track overall self-motion, while remaining partially separable at the population level. Beyond these rapid responses, passive motion also drives recurrent dynamics that persist for several seconds, thereby maintaining a temporally smooth representation.

These findings suggest that the primary visual thalamus contributes more extensively to the computation of self-motion signals than traditionally appreciated.

The ability to track both active and passive motion is required to perform robust source separation by distinguishing visual signals caused by movements of the subject from those caused by changes in the visual scene independent from subject motion. Studies in primates have shown that higher visual cortical regions can combine visual and vestibular signals to dissociate self-motion from object motion using passively imposed motion paradigms [35, 40]. A recent study in mice has also shown integration of active and passive motion signals in V1 [21]. Thus, the representations of active and passive motion in dLGN discovered in this study may provide a substrate from which downstream visual regions construct more refined estimates of self-motion and environmental motion.

The ability to separate active and passive motion signals is required to distinguish retinal signals caused by active movements from those caused by external forces acting on the subject (e.g. gravity or other external forces). Such computations may contribute to behaviours that depend on estimating the causes of self-motion, including postural control and balance, known to also depends on vision [41-43]. Here we discovered that partial separation is already present at the level of dLGN. It remains unclear to what extent downstream visual cortical areas can distinguish active from passive components of self-motion. The great diversity of movement-related signals reported in visual cortex [26, 44] and other cortical regions [44, 45] raises the possibility that active and passive motion may become increasingly differentiated across successive stages of visual processing. However, the extent to which the representation of active and passive motion is separable in V1 and other visual cortices remains unknown.

Because motion is often correlated in time, representing its recent history may help maintain a stable estimate of self-motion. Such temporally extended representations could support visual perception and motor behaviours in complex environments, where sensory information must be integrated over multiple timescales. Here, we show that neurons in dLGN and surrounding regions maintain information about passive motion over timescales of several seconds. Our results indicate that this persistence is supported by recurrent dynamics engaged by passive motion signals. They also suggest that under deep anaesthesia the substantial reduction in responses to motion could be accounted by reduced recurrent dynamics. How such recurrent dynamics arise remains unclear. One possible substrate is the thalamocortical loop linking dLGN and V1. Indeed, infragranular neurons in V1 that project to dLGN have been shown to encode passive head movements [20-22]. However, such thalamocortical loops are likely more extended as passive head movements evoked widespread cortical responses beyond V1 [21].

The thalamus has been proposed to act as a high-dimensional switchboard that flexibly gates cortical computations by modulating flow of sensory information and selectively routing such information to different cortical populations [46, 47]. The ability of dLGN to simultaneously encode active and passive motion while preserving partial separation between them is consistent with this view. Rather than fully resolving the distinction between active and passive motion, dLGN may provide a partially disentangled representation that allows downstream cortical circuits to selectively access different motion components. Such routing could facilitate subsequent computations, including the refinement of active-passive motion separation or the integration of both signals into a unified estimate of self-motion.

How cortical circuits exploit dLGN representations of active and passive motion remains an open question. Future studies combining cell-type-specific recordings, causal manipulations and naturalistic behaviour will be needed to determine how active and passive motion signals are transformed across the visual hierarchy. Such work may provide new insights into the neural mechanisms underlying the perception of self-motion, the maintenance of visual stability in dynamic environments, and the visual computations that contribute to balance and postural control.

## Methods

### Animals

Experiments were conducted on 21 adult C57BL/6J mice (19 males, 2 females) from our local colony in Manchester. All mice were initially stored in cages of 5 individuals and housed individually after surgical implantation of the chronic electrode. Animals were provided with food and water ad libitum throughout their life and kept on a 12:12 light dark cycle.

### Ethical statement

Experiments were conducted in accordance with the Animals, Scientific Procedures Act of 1986 (United Kingdom) and approved by the University of Manchester ethical review committee.

### Surgical Procedures

For recovery surgery for brain implants mice were anaesthetised with isoflurane (flow rate: 1.5-2L/min). Concentrations of 4 – 5% and 1 – 1.5% were used respectively for induction and maintenance of anaesthesia. The level of anaesthesia was verified by the lack of withdrawal reflex. After anaesthetic induction the animal head fur was trimmed. The animal was then placed in a stereotaxic frame (Narishige, Japan) and animal’s body temperature was automatically maintained at 37 C by a heating mat while animal’s eyes were protected from drying out by eye drops. Topical application of 1% EMLA cream was followed by a surgical incision to expose the skull. Two slotted cheese machine screws (M1.6x2.0mm, Precision Tools, UK) were inserted respectively into the parietal, and frontal plates to act as anchors for the dental cement and for electrical grounding of the electrode. After craniotomy the electrode was inserted into the dLGN (coordinates from bregma: 2.0-2.2mm medial-lateral, 2.3-2.5 mm rostro-caudal) at a depth of 4-4.5mm from the brain surface. Before brain insertion, electrodes were painted with a fluorescent dye to enable postmortem histological localisation. To assess electrode placement, we monitored light responses during surgical implantation of the electrode. Light-curing cement (X-tra base, VO64434-A, VOCO) was applied to seal the implant. After surgery, the mouse was released from the ear bars and allowed to recover in a single-housed heated cage. Analgesia was provided with a subcutaneous injection of 0.05mg/kg buprenorphine. After the procedure the animal was allowed to recover in a single-housed home-cage for a minimum of six days prior to experimentation. After all data were collected animals were sacrificed and histological post-mortem anatomy was performed to confirm electrode placement.

The same recovery procedures (including anaesthesia and analgesia) were used for mice in which we performed eye tracking. In these animals the base of the eye tracker implant was anchored to the skull by using 3 screws, two in left and right parietal plates and one in the frontal plate. Eye tracking cameras were attached to the base before each experimental session by a brief (∼2 minutes) anaesthesia and removed after the end of the session.

To record anaesthetised mice on the platform (**Supplementary Figure 2A**) we performed terminal procedures in which the animals were anaesthetised with urethane (1.5mg/kg) and sacrificed on the same day at the end of the experiment.

### Experimental Set-Ups

Animals were placed in a small open field arena (dimensions: 20cm x 20cm). The floor of the arena was suspended and attached to the shaft of a digital servo motor (SunFounder, 25Kg, DC 4.8-7.2v) to generate left and right tilts (**Figure 1A, C**). Visual stimulation was supplied by monitors mounted around the arena. The effective irradiance for BRIGHT stimuli was 2.7*1010 photons/cm2/s, 1.4*1013 photons/cm2/s, 1.7*1013 photons/cm2/s and 1.9*1013 photons/cm2/s respectively for S-cone opsins, rhodopsins, melanopsins and M-cone opsins.

Video recordings were performed with 8 cameras (Chamaleon 3 from Point Grey; frame rate = 20Hz) mounted over the arena (**Figure 1A**). Infrared cut-on filters (cut-on at 720 nm, Edmund Optics,#65-796) were mounted on each camera to remove visible light. Infrared illumination was provided by two lamps (Edmund Optics; 81mm IR 940nm, #18-607) placed on top of the arena. The direct infrared light projected on the arena was filtered with diffuser films. An illustration of these recordings in provided in **Supplementary Movie 1**. A full list of materials for the behavioural rig set-up and calibration is provided in [48].

Anaesthetised animals were recorded on a custom-made motorised platform controlled by an Arduino Uno board (arduino.cc). The platform rotation was measured using a tri-axial inertial measurement unit (IMU; MPU6050, InvenSense/TDK) mounted to the platform. Before each recording, a brief static calibration was performed to estimate and subtract gyroscope zero-rate offsets, and axis alignment verified using known manual rotations of the platform. Data were sampled at 80Hz. Post hoc, gyroscope readings were converted to physical units (°/s) and any drift observed during stationary periods was corrected for.

Extracellular brain recordings were performed with Neuropixel 1.0 electrodes (Imec) and a NI PXIe-1071 acquisition box (National Instruments) running at 30 kHz. Extracellular action potentials were firstly automatically sorted with Kilosort [40], then manually curated with Phy [41]. Single unit responses to light and tilt were measured as the distribution of differences between baseline and evoked spike counts across trials. Responses were identified as statistically significant when the p-value of a nonparametric sign-test applied the response distribution was lower than 0.05.

The acquisition of neural data was synchronised with the video frame acquisition via TTL pulses driven by an Arduino Uno board (arduino.cc). The same board was also used to control the stepper motor to drive the arena tilts. Psychopy software ( [42], version 2022.2.5) was used to trigger the Arduino Uno board and generate the visual stimulation.

### Reconstruction of 3D poses

We trained Deeplabcut [48] on ∼2600 manually labelled frames and used to track animals’ and arena 2D keypoint coordinates from individual cameras. Calibration for estimating camera matrices are described in [49]. These 2D coordinates were triangulated using the direct linear transformation algorithm to estimate animals’ 3D poses and arena 3D coordinates. A statistical shape model (SSM, [23, 24]) was then estimated on a subset 7000 poses from 7 mice. We used the SSM to fit the 3D poses on all animals, correct outliers and fill up missing data as described in [2].

### Separation of Active and Passive Motion

The 3D animal’s poses can be expressed as:

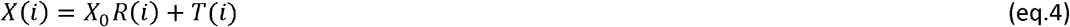

Where: *X*(*i*) is an *N*_*x*_ x 3 matrix representing respectively the 3D pose of mouse at *i*^th^ frame (*N*_*x*_ is the number of mouse keypoints); *X*_0_ represents the animal’s “reference” pose in the centre of the static arena; *R*(*i*) and *T*(*i*)are respectively 3 x 3 rotation and *N*_*x*_ x 3 translation matrices (**Figure 3A, OBSERVED MOTION**). We represent the effect of active and imposed motion at each frame as a sequential composition of two rigid transformations – the first expressing animal position from a frame aligned to the platform the second expressing transformation between such reference and a fixed external reference frame (**Figure 3A, REFERENCE FRAMES**). According to this approach, eq.4 can be further decomposed as:

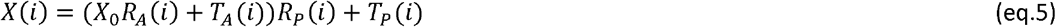

Where *R*_*A*_(*i*)and *T*_*A*_(*i*) are rotations and translations matrices associated with active motion, while *R*_*P*_(*i*) and *T*_*P*_(*i* )are associated with passive motion. For simplicity we transformed all 3D arena and mouse coordinates so that *R*_*P*_(*i*) = *I*_3_ (identity matrix) and *T*_*P*_(*i*) = 0_*Nx x* 3_ (zero matrix) when the arena is “at rest” before each tilt. The terms *R*(*i*), *T*(*i*), *R*_*P*_(*i*) and *T*_*P*_(*i*) are already given by the 3D reconstruction of the mouse and the arena coordinates. Therefore, we can combine eq.4 and eq.5 to estimate rotation and translation associated with active motion as:

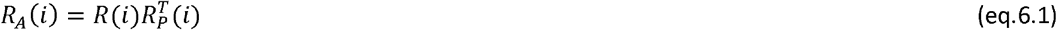

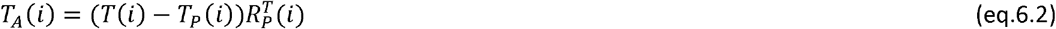

Where the superscript *T* indicates a transpose operation. We then applied *R*_*A*_(*i*) and *T*_*A*_(*i*) to the animal’s reference pose:

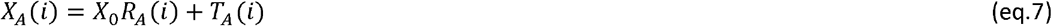

The pose *X*_*A*_(*i*) represents the animal’s pose after “undoing” the rotation and the translation imposed by the tilt (**Figure 3A, ACTIVE MOTION**). Therefore, *X*_*A*_(*i*) is solely determined by active motion. This corresponds to expressing animal’s position in respect to a reference frame continuously updated to keep it aligned to the platform. A comparison between *X*_*A*_(*i*) and *X*(*i*) is illustrated in **Supplementary Movie 2**.

To calculate imposed motion, we first determined the arena’s rotation and translation between two consecutive frames as:

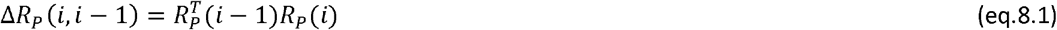

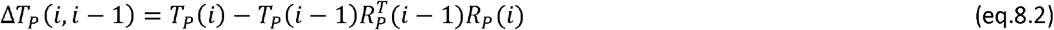

We used these transformations to determine the animal’s passive pose at *i*^th^ frame as:

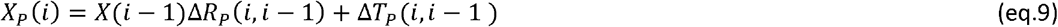

Note that, by using to this definition, we “freeze” mouse pose *X*(*i*− 1) between two consecutive frames (**Figure 3A, PASSIVE MOTION**). In this way, the only motion accounted for is that imposed by the tilt.

To calculate the linear and angular active motion each pair of consecutive poses *X*_*A*_(*i* − 1) and *X*_*A*_(*i*) we applied Procrustes superimposition to calculate the rotation 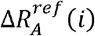 and translation 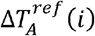 between the pose pair. Finally, the magnitude of linear active motion and the angular active motion were calculated as:

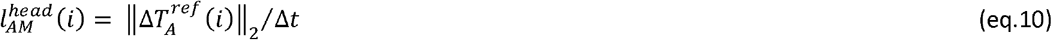

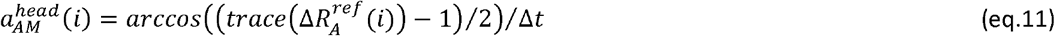

Where: ∥ · ∥ indicates the Euclidean norm and *trace*(·) indicates the trace operator. The angular active motion components associated with egocentric *x,y* and *z* axes were extracted from the rotation matrix 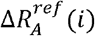 according to the rotation order *Z,Y,X* (MATLAB function: rot2eul).

To extract linear and angular passive motion we applied the same calculations described for the active case, but we used the poses *X*(*i*− 1) and *X*_*P*_(*i*) instead of *X*_*A*_(*i*− 1) and *X*_*A*_(*i*).

All motion calculations were then reapplied to the body (excluding head coordinates) in the same way.

### Eye Tracking

Miniaturised camera sensors (OV9734, Omnivision) were used to video animals’ eyes. The sensors were installed in customized tethered acquisition system (Shenzhen Yuweizhixin Electronics Co., Ltd.) equipped with an infrared cut-on filter and infrared LEDs. Frame acquisition for eye tracking was synchronised with frame acquisition from head and body tracking cameras by using Psychopy (version 2022.2.5). An illustration of these experiments is provided in **Supplementary Movie 2**.

Tracking of the eye keypoints was performed with Deeplabcut [22]. The measurements of pupil radius and eye movements on the video were first normalised by the distance between inner and the outer canthi, to correct for slight variations in camera angles across the two eyes and across animals.

Head and body 3D reconstruction and motion calculations followed the same pipeline described in the previous sections (see **Reconstruction of 3D poses** and **Separation of Active and Passive Motion**).

### Clustering Analysis

Single cell PSTHs were first z-score normalised. The clustering was performed by using a community detection algorithm [45] as implemented in the brain connectivity toolbox [46]. This algorithm receives as input an undirected connection matrix and automatically determines the number of cell clusters (or classes). To estimate the connection matrix, we applied K-means across a range of predefined cluster numbers (range: 1-20). We then estimated the connection between cell pairs as the fraction of times in which the pair was assigned to the same cluster by a K-means run. Note that all connection values were bounded between 0 and 1. All clustering analyses were performed on the joint datasets of awake and anaesthetised recordings.

### Generalised Linear Model Analysis

Motion covariates, defined as *X*_*m*_ ϵ *R*^*Tx* 4*K*^, were derived from linear and angular components of active (*l*_*AM*_, *a*_*AM*_) and passive (*l*_*PM*_, *a*_*PM*_) motion. These variables, originally sampled at *dt* = 50ms, were concatenated across a temporal window of *K* samples. In this way each row of *X*_*m*_ contained the recent history of motion signals:

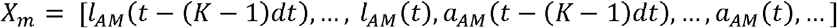

Contextual covariates, defined as *X*_*c*_, were obtained by averaging over the same temporal window and included the sine and cosine of head orientation and the distance between the animal’s head and the arena tilt axis. Visual covariates, when present, were defined as *X*_*vis*_ and derived from five variables capturing tonic and transient changes in light intensity.

Covariates associated with intrinsic fluctuations in population activity, defined as *X*_*intr*_ were derived separately for each cell *i*. Thus, by denoting the population activity *Y* ϵ *R*^*Tx N*^, we define *Y*_*i*_ ϵ *R*^*Tx* (N−1)^ as the activity of all neurons excluding cell *i*:

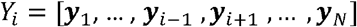

We then removed from *Y*_*i*_ the components linearly explained by all other covariates *M* = [*X*_*m*_, *X*_*c*_, *X*_*vis*_] by constructing the projection matrix *P*_*M*_ = *M*(*M*^*T*^*M*)^−1^ *M*^*T*^ and computing the residual activity 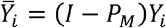. A Principal Component Analysis (PCA) was then performed on 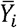 to extract the first 3 PCs, denoted as *V*. Intrinsic covariates were finally defined as 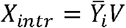. To avoid leakage between training and predictions, both *P*_*M*_ and *V* were estimated on the training set while prediction accuracy was evaluated on held out data.

Since columns of *X*_*M*_ were correlated in time and across active and passive components, we explored alternative decorrelated representations of motion covariates using both principal component analysis (PCA) and reconstruction independent component analysis (ICA, [50]). PCA and ICA were applied separately to *X*_*m*_ yielding two alternative representations *PC* = *X*_*m*_,*W*_^*PC*^_ and *IC* = *X*_*m*_,*W*_*IC*_, where *W*_*PC*_ and *W*_*IC*_ denote respectively the PCA and ICA transformation matrices. The number of PCs was set to capture 95% variance of the original signals, and we used a matching number of ICs. The design matrices for principal and independent component transformations were then defined as:

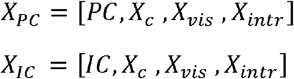

To fit spike counts from individual cells we used a Poisson GLM regression with a non-canonical “Softplus” link function expressed as ln(1 + *e* ^*βX*^) to avoid unrealistically large firing-rate predictions. We first applied lasso regularisation (MATLAB function: lassoglm) to minimise the following loss:

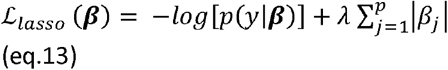

Where: ***β*** = (*β*_1_, …, *β*_*p*_) represents the parameters of the model. The regularisation parameter *λ* was determined for each cell using a five-fold cross-validation by selecting the most regularised model whose performance was within one standard error of the minimum deviance. This penalised regression was used to select the non-zero covariates. A non-penalised regression was then applied to this reduced set of covariates and model accuracy was measured on held-out data.

The model parameters were estimated on 50% data while the remaining 50% was used to evaluate goodness of fit. To avoid leakage between training and performance evaluation, training and held out samples were divided according to blocks defined by separate experimental trials. To measure the goodness-of-fit we first calculated residual and null deviances defined as:

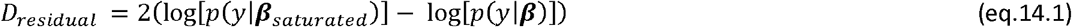

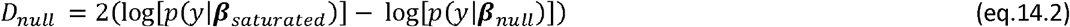

Where: ***λ***_*saturated*_ represents the parameters of the saturated model in which each observation is fitted exactly and ***λ***_*null*_ the parameters of the model with the intercept only. We used *eq*.*14*.*1* and *eq*.*14*.*2* to calculate a pseudo *R*^2^ defined as:

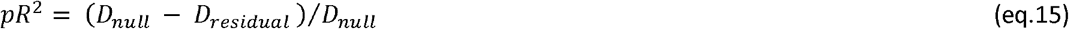

This measure, equivalent to *R*^2^ in least square estimates, quantifies deviance reduction for maximum likelihood estimates [51].

### Population Decoding Analysis

We applied a linear decoder to all simultaneously recorded cells that responded to tilt and light. Estimation of the linear decoder parameters was regularised with ridge regression (MATLAB function: ridge) to minimise the following cost:

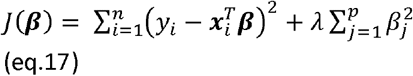

Where: ***β*** = (*β*_1_, …, *β*_*p*_) represents the parameters of the model, 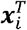 the the vector of spike counts at *i*^*th*^ frame, *y*_*i*_ the motion speed signal to be decoded. The ridge parameter variable *λ* was optimised once across all datasets that we used for this analysis. A 50% of the data were held out and used to evaluate the decoding performances.

### Separation of rapid responses and recurrent dynamics

To separate the neural influence of passive motion signals on rapid responses and recurrent dynamics we used the IPSID algorithm [25], that defines a dynamical system as:

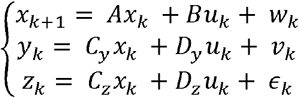

Where 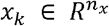 represents the latent dynamical state, 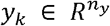 the neural activity, 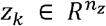 the behavioural signals, 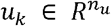 the external input to the system, the terms *W*_*k*_, *V*_*k*_, *ϵ*_*k*_ Gaussian white noise (see for [25] details).

To apply IPSID to our data we defined *z*_*k*_ = [*l*_*AM*_ (*k*), *a*_*AM*_ (*k*)] and *u*_*k*_ = [*l*_*PM*_ (*k*), *a*_*PM*_ (*k*)], so that passive motion signals acted as external inputs driving both active-motion behaviour and neural activity. For data recorded during visual stimulation we also included an additional input associated with light intensity. Neural activity comprised all the tilt- and light-responsive neurons simultaneously recorded from each animal. We first estimated the parameters of such model through five-fold cross-validation by using the Python package PyPSID and applying a trial-wise five-fold cross-validation procedure. Model parameters were estimated independently from the training trials of each fold and held-out trials were used for dimensionality selection and performance evaluation.

Within the IPSID framework the *n*_*x*_ dimensions of the latent dynamical state can be further decomposed into *n*_1_ dimensions driving both behaviour and neural activity, *n*_2_ dimensions driving only neuronal activity and *n*_3_ dimensions driving only behaviour. Following this order, we sequentially selected *n*_1_, *n*_2_, and *n*_3_ through a systematic grid search using the one-standard-error rule. This procedure selected *n*_1_=1, *n*_2_=9, and *n*_3_ =0, indicating that additional behaviour-specific dimensions contained in *n*_3_ were not required for our datasets. Prediction performances were measured through Pearson’s correlation coefficient. This that was computed within each fold after concatenating samples from independently predicted held-out trials and averaged across behavioural dimensions (for estimating *n*_1_ and *n*_3_) and cells (for estimating *n*_2_).

To separate direct and recurrent drives on neuronal activity, defined respectively by projections *D*_*y*_*u*_*k*_ and *C*_*x*_*X*_*k*_, we simulated the fitted IPSID model with no additional noise and separately ablated each component. Through these simulations we obtained two instances of population activity *y*_*k,direct*_ and *y*_*k,recurrent*_. We then used the equivalence *Var*(*y*_*k*_) = *Var*(*y*_*k, direct*_) + *Var*(*y*_*k, recurrent*_.) +2 *COV*(*y*_*k, recurrent*_) to estimate the relative contribution of each component to neural activity.

To estimate the timescales over which recurrent dynamics influence neural activity we first decomposed the recurrent matrix as A = *VΛV*^−1^ where the columns of *V* correspond to eigenvectors and the diagonal elements of Λ the associated eigenvalues. We then computed observations matrix *C*_*y*._ Each element of *M* quantifies the weight assigned by each neuron to a given latent dynamical component. The time constant associated with each eigenvalue *λ*_*i*_ was defined as *τ*_*i*_ = − Δ*t/*log(|*λ*_*i*_|) . For each neuron, time constants were grouped into logarithmically spaced intervals, and the amplitudes of the corresponding projections, defined by *M*, were accumulated within each interval to quantify the representation of latent dynamics across timescales.

## Declaration of generative AI and AI-assisted technologies in the manuscript preparation process

Generative AI and AI-assisted technologies were used solely to improve the language, grammar, and clarity of the manuscript. The authors reviewed and edited all generated text and take full responsibility for the content of the publication.

## Acknowledgements

We wish to express thank Robert J Lucas (R.J.L), Tim Brown and Annette Allen for intellectual and technical support throughout this study. Annette Allen also provided key expertise in developing some of the illustrations for this study. This study was funded by Sir Henry Dale fellowship from Wellcome Trust to R.S. (220163/Z/20/Z); a research grant by Biotechnology and Biological Sciences Research Council to R.S.P. and R.S. (BB/V009680/1); a Wellcome Trust Investigator award to R.J.L. (210684/Z/18/Z).

## Author Contributions

R.S., A.S.E. and R.S.P designed the study; R.S., M.P.H., A.S.E., Q.H., F.J. performed the experiments; A.S.E., R.S., Z.M., F.P.M., F.J., K.S., R.J., K.T. provided source code and analysed the data. R.S., A.S.E. and R.S.P. wrote the manuscript.

## Declaration of Interests

The authors declare no competing interests.

## Supplementary Results

We asked whether responses to tilt events comparable to those observed in awake, freely moving animals also occur in anaesthetised animals, in which active eye, head and body movements are abolished. After the induction of a surgical level of anaesthesia (n = 7 animals, 1.5 mg/kg of urethane), animals were placed on a motorised platform designed to deliver tilts along the pitch and roll axes (**Supplementary Figure 1A, B**). An LED, stably mounted on the platform, enabled us to deliver the same visual stimulation when the platform was stationary or tilting.

The dataset of visual responses in absence of tilts in anaesthetised animals was merged with data from awake animals. This enabled us to classify light responses in anaesthetised conditions along the same five classes described in awake animals (**Supplementary Figure 1C, E, left panel; Supplementary Figure 1D, F, orange lines**). Compared with awake animals, we found that, under anaesthesia, visual responses were only minimally impacted by tilt (**Supplementary Figure 1C, E, mid panel; Supplementary Figure 1D, F, blue lines**). Consistent with this, the amplitude of tilt responses in the dark were negligible when compared with light responses (**Supplementary Figure 1C, E, right panel; Supplementary Figure 1D, F, black lines**). The effect of tilt on light responses was also qualitatively different and, when significant, comprised both increments and decrements in firing rates (**Supplementary Figure 1G, H, I, J**). These results indicate that tilt events evoke qualitatively and quantitatively different responses in anaesthetised animals.

**Supplementary Figure 1.**
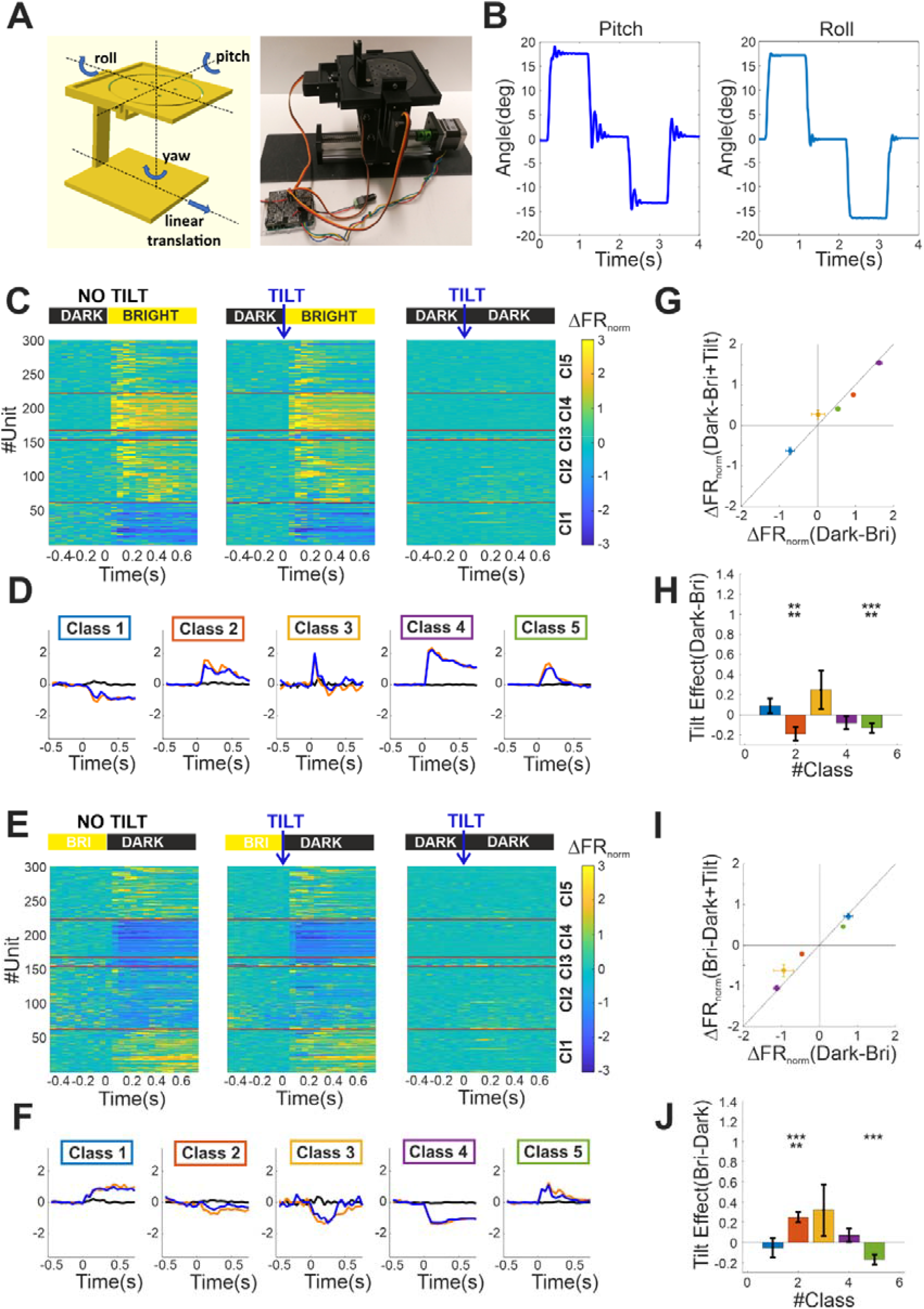
**A)** Schematic (left) and picture (right) of the motorised platform we used for delivering passive motion to anaesthetised animals. In the schematic we highlight the tilt axes. Animals are placed on the top of the arena and secured by head fixation. **B)** Gyroscopic measures of the pitch and roll angles during passive motion stimulation. **C-I)** Same a **Figure 2A-H** but here we report data collected in anaesthetised animals mounted on the platform shown in panel **A**.

## Notes

https://doi.org/10.17605/OSF.IO/DRVFP

### Competing Interest Statement

The authors have declared no competing interest.

### Summary of Updates

The study has been extended, thoroughly re-analyzed and extensively rewritten. The current study includes a novel framework for separating active and passive motion, new population-level decoding analyses demonstrating their partial separability, and dynamical systems modelling showing temporally extended representations. All these additions substantially strengthen and expand the scope of the work.

